# AMP-activated protein kinase is a key regulator of acute neurovascular permeability

**DOI:** 10.1101/802033

**Authors:** Silvia Dragoni, Bruna Caridi, Eleni Karatsai, Thomas Burgoyne, Mosharraf H. Sarker, Patric Turowski

**Affiliations:** Institute of Ophthalmology, University College London, 11-43 Bath Street, London EC1V 9EL, UK; NIHR Moorfields Biomedical Research Centre, Moorfields Eye Hospital, London, UK; School of Science, Engineering & Design, Teesside University, Stephenson Street, Middlesbrough TS1 3BA, UK

**Keywords:** Neurovascular leakage, AMP-activated protein kinase, VEGF-A, bradykinin, retina

## Abstract

Many neuronal and retinal disorders are associated with pathological hyperpermeability of the microvasculature. We have used explants of rodent retinae to measure and manipulate acute neurovascular permeability and signal transduction to study the role of AMP-activated protein kinase (AMPK). Following stimulation with either vascular endothelial growth factor (VEGF-A) or bradykinin (BK), AMPK was rapidly and strongly phosphorylated and acted as a key mediator of permeability downstream of Ca^2+^ and Ca^2+^/calmodulin-dependent protein kinase kinase (CAMKK). Accordingly, AMPK agonists potently induced acute retinal vascular leakage. AMPK activation led to phosphorylation of endothelial nitric oxide synthase (eNOS), which in turn increased VE-cadherin phosphorylation on Y685. In parallel, AMPK also mediated phosphorylation of p38 MAP kinase and HSP27, indicating that it regulated paracellular junctions and cellular contractility, both previously associated with endothelial permeability. Endothelial AMPK provided a missing link in neurovascular permeability, connecting Ca^2+^ transients to the activation of eNOS and p38, irrespective of the permeability-inducing factor used. Collectively, the ex-vivo retina model was easily accessible and, due to its compatibility with small molecule antagonists/agonists and siRNA, constitutes a reliable tool to study regulators and mechanism of neurovascular permeability.

## INTRODUCTION

Leakage within the vascular system can be the cause or a significant co-morbidity of a variety of pathologies and is the consequence of endothelial hyperpermeability, which leads to extravasation of fluids and proteins, resulting in interstitial oedema^1^. In the nervous system, where the vasculature is uniquely impermeable and is referred to as the blood-brain barrier (BBB), vascular leakage accompanies stroke, multiple sclerosis, Parkinsons Disease as well as various forms of dementia^2, 3^ Neurovascular leakage also affects the blood-retinal barrier (BRB), where it is a hallmark feature of diabetic retinopathy and neovascular age-related macular degeneration^4^. Leakage during retinopathies is driven by permeability-inducing factors (PIFs), most prominently by the angiogenic growth factor VEGF-A; and VEGF-A antagonists are successfully used to reduce oedema and abnormal vessel growth, and restore neuronal dysfunction^5^. Meta-analysis of the clinical use of anti-VEGFs in diabetic macular oedema suggests that PIFs other than VEGF-A play an important role in the pathogenesis of retinal leakage disease^6^, including angiopoeitin-2^7^, lysophosphatidylcholine^8^ and BK^9^.

PIFs induce both acute and chronic vascular leakage. For instance, exposure of the vascular endothelium to VEGF-A leads to acute permeability that usually lasts less than 30 minutes. If not resolved persistent leakage ensues that chronically impairs vascular integrity^1, 10^ Different PIFs bind to distinct endothelial cell (EC) surface receptors, but ultimately all permeability responses involve paracellular junction modulation or formation of transport vesicles^1, 2^, suggesting that ECs regulate hyperpermeability through a core molecular machinery and common downstream signalling. Indeed, e.g. Ca^2+^ transients, phosphorylation of the MAP kinase p38 and enhanced actin contractility are associated with all vascular permeability, as is activation of eNOS^11,12,13,14^. Adherens and tight junction modulation is associated with paracellular permeability and the phosphorylation of VE-cadherin (VE-cad) is associated with acute permeability in the periphery and the retina^15, 16^ In the retina the phosphorylation of occludin on S490 and its subsequent internalisation also plays an important role, at least in a more chronic setting^17^. Identifying core signalling, on which all leakage responses depend, is desirable for therapeutic development as it would allow treatment without prior knowledge of specific known (or unknown) extracellular PIF.

In the brain and the retina, entirely different signalling is induced by VEGF-A in ECs when it is added luminally (from the blood side) or abluminally (from the tissue side), with leakage-inducing signalling entirely restricted to abluminal stimulation^12^. For instance, leakage-associated p38 activation is triggered by abluminal (basal) VEGF-A stimulation, whilst activation of the PI3K/AKT pathway, which does not mediate permeability, is only seen following luminal (apical) stimulation. Thus, signalling specific to leakage can be inferred by comparing cellular stimulation following luminal and abluminal addition of VEGF-A. Conversely, BK efficiently induces permeability from the basal as well as apical side of cerebral or retinal ECs.

AMP-activated protein kinase (AMPK) is a phylogenetically conserved energy sensor that regulates energy homeostasis by coordinating metabolic pathways and thus balancing energy requirement with nutrient supply^18^. Previous studies suggest that AMPK acts as a protector of the BBB integrity, for instance by preventing LPS-enhanced NAD(P)H oxidase expression in ECs and the consequent barrier dysfunction and enhanced permeability^19^. Moreover, AMPK mediates up-regulation of BBB functions induced in vitro by metformin, a drug used for the treatment of diabetes^20^. Nevertheless, in the retina AMPK activation can lead to the breakdown of the outer, non-vascular BRB, which is constituted by the retinal pigment epithelium^21^. However, if and how AMPK contributes to permeability induction by agonists such as VEGF-A or BK in neural microvessels is unknown.

To identify and validate core components mediating acute permeability in neurovascular ECs, we adopted an ex vivo retinal preparation, originally described for rats^22^. Development of this method allowed measurement of real time changes of permeability and signalling in intact BRB vessels from rat and mouse. Importantly, this model system was compatible with precise pharmacokinetic agonist studies, parallel IHC staining and manipulation using siRNA. Our workflow can be used to identify core regulators of CNS endothelial hyperpermeability and was validated by identifying AMPK as a novel, key regulator linking VEGF-A or BK-induced Ca^2+^ transients to eNOS activation and VE-cad phosphorylation.

## MATERIALS AND METHODS

### Materials

Recombinant rat VEGF-A (165) was purchased from R&D Systems (Abingdon, United Kingdom). Bradykinin, Sulforhodamine-B, Evans blue, SU-1498, SB-203580, Compound-C, STO-609, L-NAME, BAPTA-AM, AICAR and A769662 were purchased from MERCK (Dorset, United Kingdom). Polyclonal antibodies specific for p38, Hsp27, AMPKα, eNOS and their phosphorylated forms (p38 Thr180/Tyr182, pHSP27 Ser82, AMPK Thr172 and eNOS Ser1177) were from Cell Signaling Technology (Beverly, MA). Polyclonal antibodies against phosphorylated VE-cad (p-Y658-VEC and p-Y685-VEC) were a gift from E. Dejana (Milan, Italy).

### Animals

Wistar female rats (7-10 weeks old) and C75BL/6J mice (7-12 weeks old) were purchased from Charles River Laboratories. All animal procedures were performed in accordance with Animal Welfare Ethical Review Body (AWERB) and Association for Research in Vision and Ophthalmology (ARVO) Statement for the Use of Animals in Ophthalmic and Vision Research guidelines and under a UK Home Office licence.

## Methods

### Brain microvascular EC isolation and culture

Microvessels were isolated from rat cortical grey matter by collagenase dispase digestion and BSA and Percoll density gradient centrifugation^12^. Purified vessels were seeded onto collagen IV/fibronectin-coated Costar Transwells (Fisher Scientific) at high density (vessels from 6 rat brains per 40 cm^2^). Cells were grown in EGM2-MV (Lonza), with 5 mg/ml puromycin from the second day for 3 days, for 2–3 weeks until their TEER plateaued at values above 200 Ω.cm^2^.

The human brain MVEC line hCMEC/D3 was also grown in EGM2-MV as previously described^23^.

### Transendothelial Electrical Resistance (TEER)

Changes in the TEER were determined by time-resolved impedance spectroscopy of primary cerebral rat brain microvascular ECs grown on 12 mm Transwells, using a CellZscope (Nanoanalytics). Before the addition of VEGF-A and BK, TEER values were 500-800 Ω.cm^2^.

### Immunocytochemistry

Primary cerebral rat brain microvascular ECs were fixed with methanol (−20°C). Staining was performed as previously described using antibodies against occludin (OC-3F10, Invitrogen)^24^ or VE-cad^25^.

### Immunogold Electron Microscopy

Cryo-immuno electron microscopy (EM) was performed as previously described^12^. Briefly, hCMEC/D3 cells were fixed in 4% PFA and 0.1% glutaraldehyde. Sections were stained using antibodies against the extracellular domain (TEA 1.31; Serotech) or the C terminus (sc-6458; Santa Cruz) of VE-cad. Image J (NIH) was used to process images and measure the distance among the gold particles and the interendothelial junctions.

### Retinal explant preparation

Retinal explants were prepared essentially as described before^22^. A Wistar female rats or C75BL/6J mouse was killed by overdose of CO2. The common carotid artery was carefully exposed and cannulated with a glass microcannula. The head vasculature was then flushed first with heparinized saline (300U/L heparin in 0.9% NaCl. Mouse: 5 ml; rat: 20 ml), then with stabilizing solution (10 mM Mg^2+^, 110 mM NaCl, 8 mMKCl, 10 mM HEPES, 1 mM CaCl2, pH 7.0. 10 μM Isoproterenol to be added before use. Mouse: 5 ml; rat: 20 ml), also referred to as cardioplegic solution containing isoproterenol, and finally with the same solution supplemented with 5 g/L Evans blue dye in 10% albumin (mouse: 5 ml; rat: 20 ml) for subsequent visualisation of the vasculature. Next, an eye was surgically removed and and the retina isolated, together with the attached sclera. The retina was flattened onto a transparent silicone medium (SYLGARD^®^ 184 SILICONE ELASTOMER KIT by Dow Corning), kept in position by a metal ring, and the resulting well sealed by grease. Throughout the procedure the retina was continuously superfused with Krebs solution (124 mM sodium chloride, 5 mM potassium chloride, 2 mM MgSO4, 0.125 mM NaH2 PO4, 22 mM NaHCO3, 2 mM CaCl2, pH 7.4. 5 mM glucose and 0.1% BSA to be added before use).

### Permeability measurements

Retinal explants were mounted for visualisation and further experimentation on an upright Zeiss Axiophot fluorescent microscope. A radial vessel of a superfused retinal explant was cannulated at ca. 150 μm from the optic nerve using a microinjection needle (tip diameter 1-5 μm, sharpened to a bevel of < 30°) and the entire retinal vasculature injected with sulforhodamine-B (1 mg/ml in Krebs solution). Illumination was switched to fluorescence and the vessels were visualized under a TRITC filter using a Olympus 40X water immersion objective. For permeability measurements a microvessel was chosen at least 200 μm away from the cannulated radial vessel. Fluorescent content of the vessel was recorded continuously by time lapse (1 frame/2 s) on a Hamamatsu CCD camera for at least 2 min. A baseline was recorded for ca. 30 s, before VEGF-A or BK (in Krebs solution) was added on the top of the retina. Time lapse series were analysed using ImageJ. Time-dependent fluorescence intensity data of the chosen vessel was derived from a square region of interest (ca. 18 x 18 pixels) (Supplemental Figure 1a and b). Fluorescence in the immediate vicinity of the microvessel was measured and subtracted from the vessel fluorescence measurements. Pixel intensity measurements were charted against time, and permeability values were computed by fitting data to the exponential equation Ct = C0*e^-kt^, where k = 4P/d and d is the diameter of the vessel^12^. The difference in permeability between pre-treatment and post-treatment resulted in the absolute permeability change associated with the treatment regimen.

### Immunohistochemistry

After dissection, retinae from rat or mouse were fixed with 4% PFA at room temperature for 1 h. After 30 min of blocking (3% Triton X-100, 1% Tween, 0.5% Bovine Serum Albumin in 2x PBS), retinae were incubated with primary antibodies against Isolectin B4 (IB-4), P-p38, P-HSP27, P-eNOS, P-AMPKα, claudin-5 and P-VE-Cadherin at 4 °C o/n. Retinae were washed and incubated with matching Alexa Fluor conjugated secondary antibodies at room temperature for 2 h. Finally, retinae were washed and mounted using Mowiol 4-88 mounting medium (Sigma). More details can be found in ref 12.

### siRNA-mediated Knockdown of claudin-5 and AMPKα

Specific SiRNA sequences targeting claudin-5 or the α subunit of AMPK were purchased from Dharmacon (Chicago, IL). Mice were anesthetised by intraperitoneal injection of 100 ul of 6% Narketan (ketamine) and 10% Dormitor (medetomidine) in sterile water. 2 μl of the siRNA (1ng/ml in sterile PBS) were injected intravitreously in the right eye under a stereomicroscope, using a Hamilton syringe with a 3 degrees Hamilton RN needle (Esslab). Two microliters of scrambled siRNA were injected into the left eye as a control. To inject, an initial puncture was made to the superior nasal sclera, at the level of the pars plana. Then, the tip of the needle was further introduced through the puncture hole with a 45-degree angle into the vitreous body. Retinae were isolated 72 h after the injection.

### Phosphoantibody array

Mature monolayers of primary, unpassaged brain microvascular ECs grown on 24 mm Costar Transwells were stimulated with VEGF-A (50 ng/ml) from the apical or basal side for 5 or 30 min. Cells from 2 Transwells were combined by lysis in 200 μL of lysis buffer and subjected to screening using Human Phospho-Kinase Array Proteome Profiler Array (R&D Systems; ARY003B) exactly according to the manufacturer’s instructions. Arrays were exposed for varying amounts of time to capture signals in the linear range and quantified by densitometric scanning and ImageJ (NIH). Signals were normalised using array-internal controls. Results were expressed as fold-differences between apical versus basal signals.

### Western Blots

Cell lysates were prepared as previously described^12^. Proteins were separated by SDS-PAGE and transferred to nitrocellulose by semidry electrotransfer. Membranes were blocked o/n and then incubated with the appropriate antibody diluted at 1:2,000. Membranes were washed three times with TBS/0.1% Tween-20 before 1h incubation with an anti-mouse or anti-rabbit HRP-conjugated IgG (GE Healthcare) at a dilution of 1:10,000 and 1:5,000, respectively. Membranes were developed using the ECL reagents (Roche) and exposed to X-ray film. Protein bands were evaluated by densitometric quantification, normalized against the amount of total protein, and GADPH or Tubulin.

### Statistics

TEER measurements of three independent cell monolayers were combined and expressed as mean ± SD. Significant differences were determined by two ways ANOVA with replication, with significance levels set at 0,05, followed by post-hoc Bonferroni’s multiple comparison test.

Densitometric quantification of four independent immunoblots were determined by changes in phosphoprotein content normalized to tubulin or GADPH/total protein loading controls, with values expressed as fold increase. Data were presented as mean ± SD. Statistics were performed using one-way ANOVA with significance levels set at 0.05, followed by post-hoc Dunnett’s tests.

Permeability measurements from at least 4 different ex vivo retinae were combined and expressed as mean ± SD. Significant differences were determined via t-test between the control and each inhibitor.

Significance levels were set to *, p < 0.05; **, 0.001 < p < 0.01, ***, p ≤ 0.001.

## RESULTS

### VEGF-A and BK-induced permeability and junctional changes in brain microvascular ECs

Treatment of primary rat brain microvascular ECs with VEGF-A or BK significantly reduced TEER, indicating that paracellular permeability was induced (Figure 1a and b). TEER dropped immediately and reached a minimum within less than 5 min after addition of either VEGF-A or BK before reverting to control levels within 1 h. Thereafter, another significant, but more modest reduction in TEER was observed, indicative of a more chronic change in cell monolayer permeability. In order to correlate TEER changes with paracellular junction breakdown, the distribution of occludin and VE-Cad was analysed in VEGF-A- and BK-stimulated primary brain microvascular ECs (Figure 1c). As judged by confocal microscopy, occludin expression and distribution remained unchanged for up to 2 h of VEGF-A or BK stimulation. VE-cad levels also remained unchanged, but a broadening of the staining was observed within 5 min of the addition of the PIF, in particular at and around tricellular junctions. In agreement, cryo-immuno–EM of hCMEC/D3 cells revealed a significant relocation of VE-cad from the junctions to the cell interior by an average distance of 55 and 66 nm following a 5-min stimulation with VEGF-A and BK, respectively (Figure 1d-g). These results showed that single addition of either VEGF-A or BK induced acute and chronic permeability, and that the acute response was accompanied by VE-cad redistribution away from the junctions.

**Figure 1.**
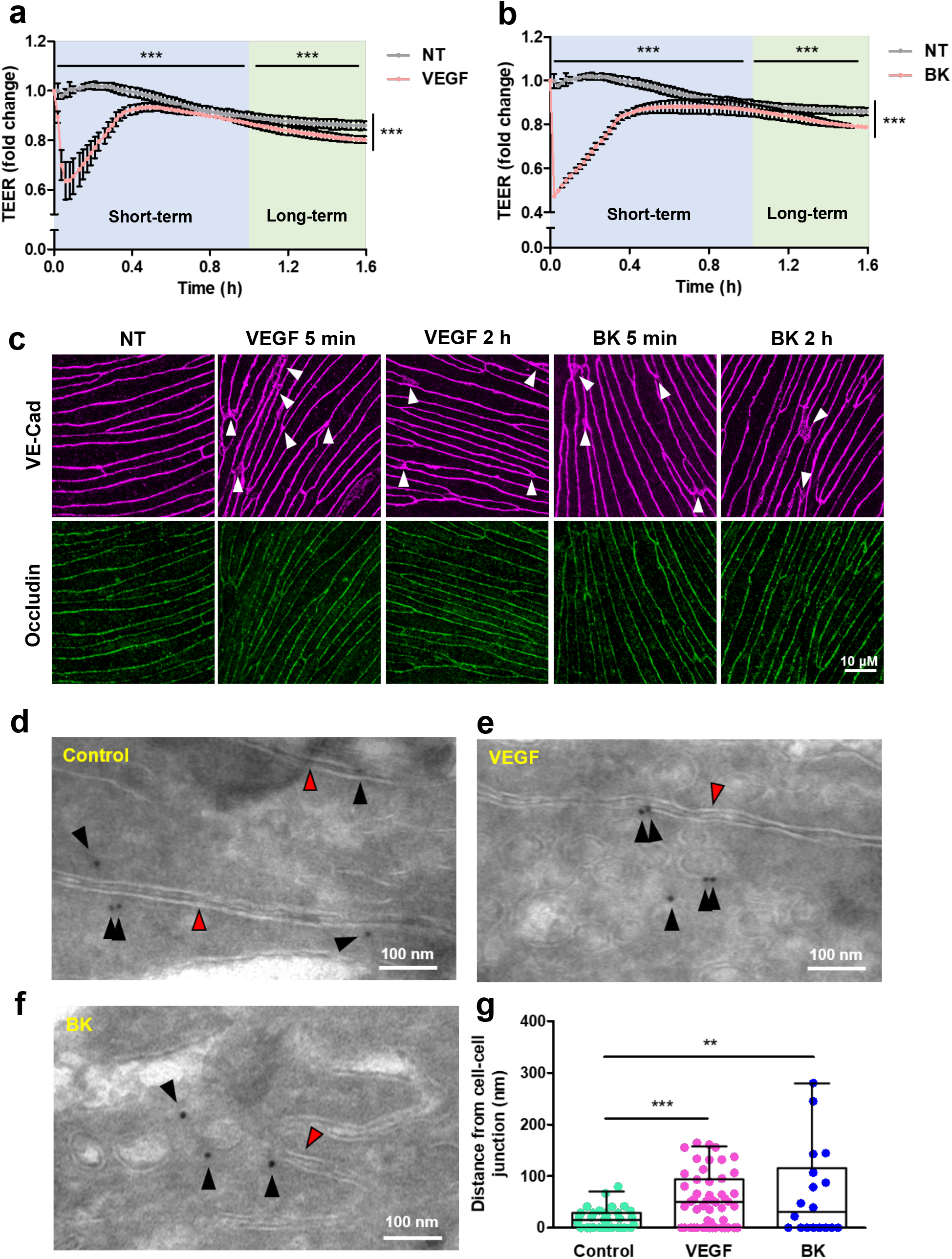
VEGF-A and BK induced permeability in cultured brain microvascular ECs. **(a, b)** Primary rat brain microvascular ECs were grown on permeable Transwell inserts to confluence and until they reached full electrical barrier (500-800 Ω.cm2). VEGF-A (a) or BK (b) were added at time 0. Shown are means ± SD of normalized resistance changes (n=3). Significant changes were detected in the short- and long-term, as well as overall responses. **(c)** Changes in the distribution of VE-Cad and Occludin in response to basal (corresponding to abluminal) stimulation with VEGF-A or BK were analysed by confocal microscopy in post-confluent primary rat brain microvascular ECs. Whites arrows indicate the broadening of the VE-Cad staining. Scale bar, 10 μM. **(d-f)** Cryo-immuno-EM of VE-Cad distribution in control (d) and VEGF-A (e) or BK (f) stimulated human hCMEC/D3 cells. Shown are interendothelial junction areas with the two adjacent membranes (red arrowheads). White arrowheads point out gold labelled VE-Cad, which in control cells was found predominantly associated with abutting plasma membranes (within 20 nm; i.e. the distance expected by the primary and the secondary bridging Ab). Scale bar, 100 nm. **(g)** Distances of VE-Cad gold particles from cell-cell junctions determined from three independent preparations as shown in (d-f). **p < 0.01, ***p < 0.001.

### Validation of a modified ex-vivo retinal model

In order to study the acute phase of VEGF-A- and BK-induced permeability in intact retinal microvessels in real time, we adopted an ex-vivo retinal model^22^. Permeability measurements used rat retinal explants, in which the vasculature was stabilised with a cardioplegic solution. To assess if the ex vivo preparation and perfusions led to alterations of the retinal vasculature and to determine stability of the preparation, we compared directly perfused fixed retinae with others, perfused with cardioplegic solution and left under superfusion with Krebs solution for 1 h before fixation. Subsequent whole mount staining for the tight junction protein claudin-5 and the adherens junction protein VE-cad revealed characteristic strands of continuous paracellular staining (Figure 2a, b) in both preparations. Importantly, the staining pattern was indistinguishable between the two different preparations, indicating that the perfusion did not cause significant disturbances of endothelial junctions. Permeability measurements were carried out by monitoring sulforhodamine B loss from individual microvessels (Supplemental Figure 1a, b). Baseline permeability to sulforhodamine-B was very low and on average 0.2 ± 0.16x 10^-6^ cm/s. Taken together these data showed that morphological and barrier properties of the retinal microvasculature were well preserved in these preparations.

**Figure 2.**
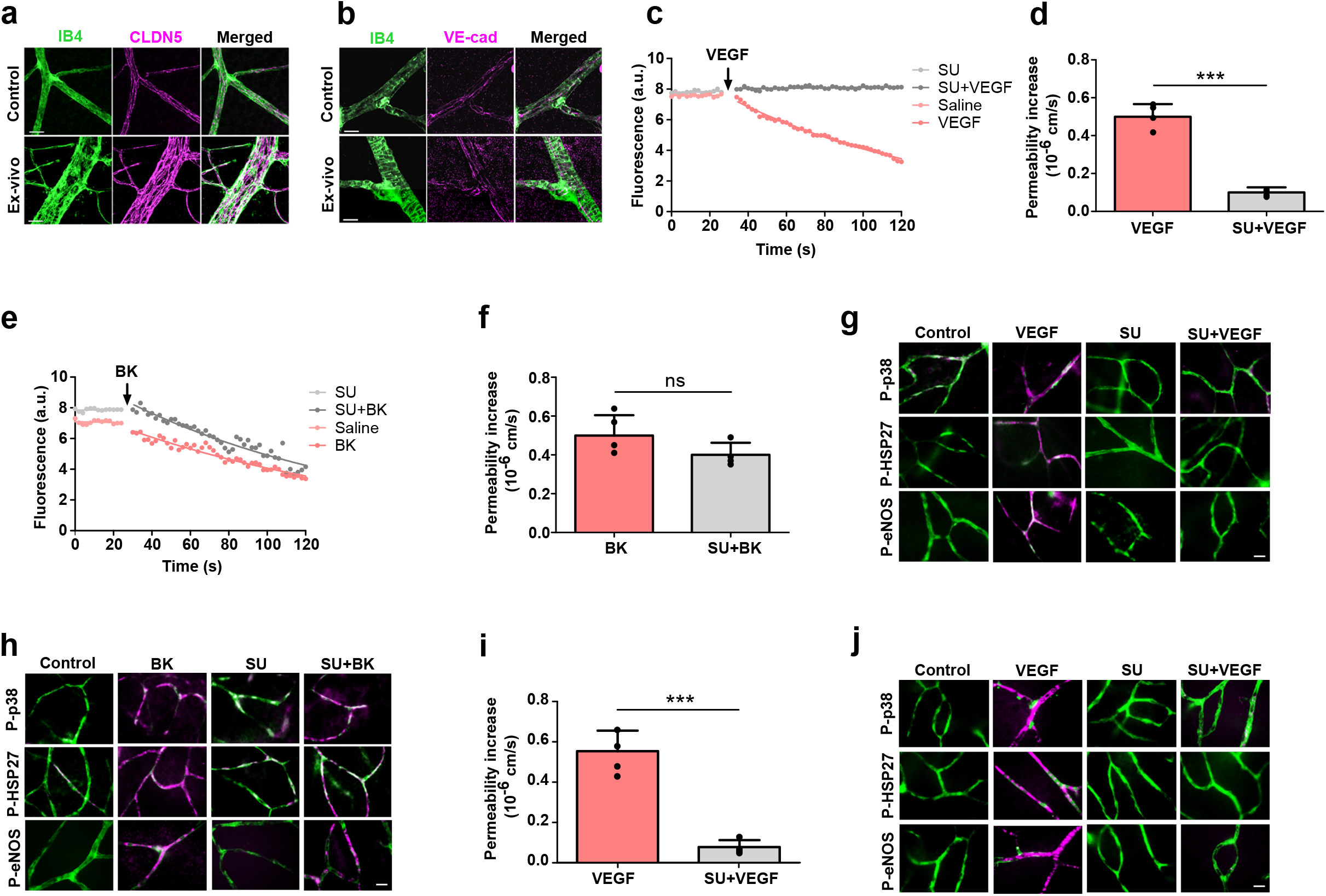
Validation of the ex vivo retina model in rats. **(a, b)** Control retinae were from animals directly perfused fixed with 4% PFA. Ex vivo retinae were isolated as described, flat mounted and left submerged with Krebs solutions for 1 h before PFA fixation. Whole mounts were stained for Isolectin B4 (IB4), claudin-5 (CLDN5) and VE-cad as indicated. **(c-f)** Sulforhodamine-B fluorescent intensities were recorded in single retinal capillaries. 50 ng/ml VEGF-A (c, d) or 10 μM Bradykinin (BK) (e, f) were applied on top of the retina (abluminal side) at times indicated. Optionally retinae were preincubated with the VEGFR2-selective antagonist SU1498 (10 μM) for 15 min prior to recording. Mean (± SD) permeability changes recorded from at least four retinae are shown in (d) and (f). **(g, h)** Ex vivo retinal explants were incubated with VEGF-A (g) or BK (h) for 2 min, fixed using 4% PFA and then stained using IB4 (green) and for phospho-p38 (pT180/Y182), phospho-Hsp27 (pS82) or phospho-eNOS (pS1177) (magenta). **(i)** Microvessel permeability changes were recorded in mouse retinae as described in (e, f). **(j)** Mouse retinal explants were stimulated and stained as described in (g) using phospho-specific antibodies to p38, HSP27 and eNOS. ns non-significant, ***p < 0.001. Scale bars, 10 μm.

Stimulation of the ex-vivo retina with VEGF-A or BK induced an immediate, marked loss of sulforhodamine B from the microvessel lumen, which was similar for both stimuli and amounted to a ca. 3-fold increase in microvessel permeability (Figures 2c-f). Preincubation of the ex-vivo retina with VEGFR2 inhibitor SU-1498 for 15 min prevented VEGF-A-but not BK-induced permeability, confirming the role of VEGFR2 in VEGF-A-induced permeability and indicating that BK acted through a different receptor. Whole mount retinal staining showed that VEGF-A- and BK-induced permeability coincided with the phosphorylation of p38 on T180/Y182, its downstream effector HSP27 (on S82), as well as eNOS (on S1177) (Figure 2g, h). Phosphorylation of all three downstream effectors in response to VEGF-A but not BK was abolished following preincubation with SU-1498.

This experimental model was also used for mouse retinae. Baseline permeability in mouse preparations was 0.15 ± 0.1×10^-6^ cm/s. VEGF-A stimulation increased permeability to 0.65 ± 0.2×10^-6^ cm/s, and this was again sensitive to SU-1498 (Figure 2i). Furthermore, we observed SU-1498-sensitive phosphorylation of p38, HSP27 and eNOS in mouse retinal microvessels within 2 min of VEGF-A stimulation (Figure 2j). To assess the compatibility of the ex vivo retina with knockdown technology, mouse eyes were injected intravitreously with siRNA against CLDN5. Western blot analysis of retinal lysates, harvested 72 h after the injection, showed that CLDN5 expression was significantly reduced by 65% (Supplemental Figure 1c, d) and this was corroborated by whole mount staining of the retina (Supplemental Figure 1e). Microvessel permeability following knock-down of CLDN5 increased ca. 3-fold (to 0.41±0.03 cm/s) (Supplemental Figure 1f, g). Taken together, these results demonstrated that the ex-vivo retinal preparation was a reliable model to measure retinal paracellular microvessel permeability in rats and mice.

### VEGF-A and Bradykinin induce AMPK phosphorylation

In order to find new regulators of permeability, primary rat brain microvascular ECs were stimulated for 5 or 30 min with VEGF-A (50 ng/ml) from either the apical (non-permeability inducing) or basal (permeability-inducing) side (Figure 3a) and cell lysates analysed by a phospho-protein antibody array. In response to VEGF-A stimulation many signalling components were phosphorylated, as exemplified by p38, HSP27, AMPK, eNOS, SRC, ERK and AKT (Figure 3b, c).

**Figure 3.**
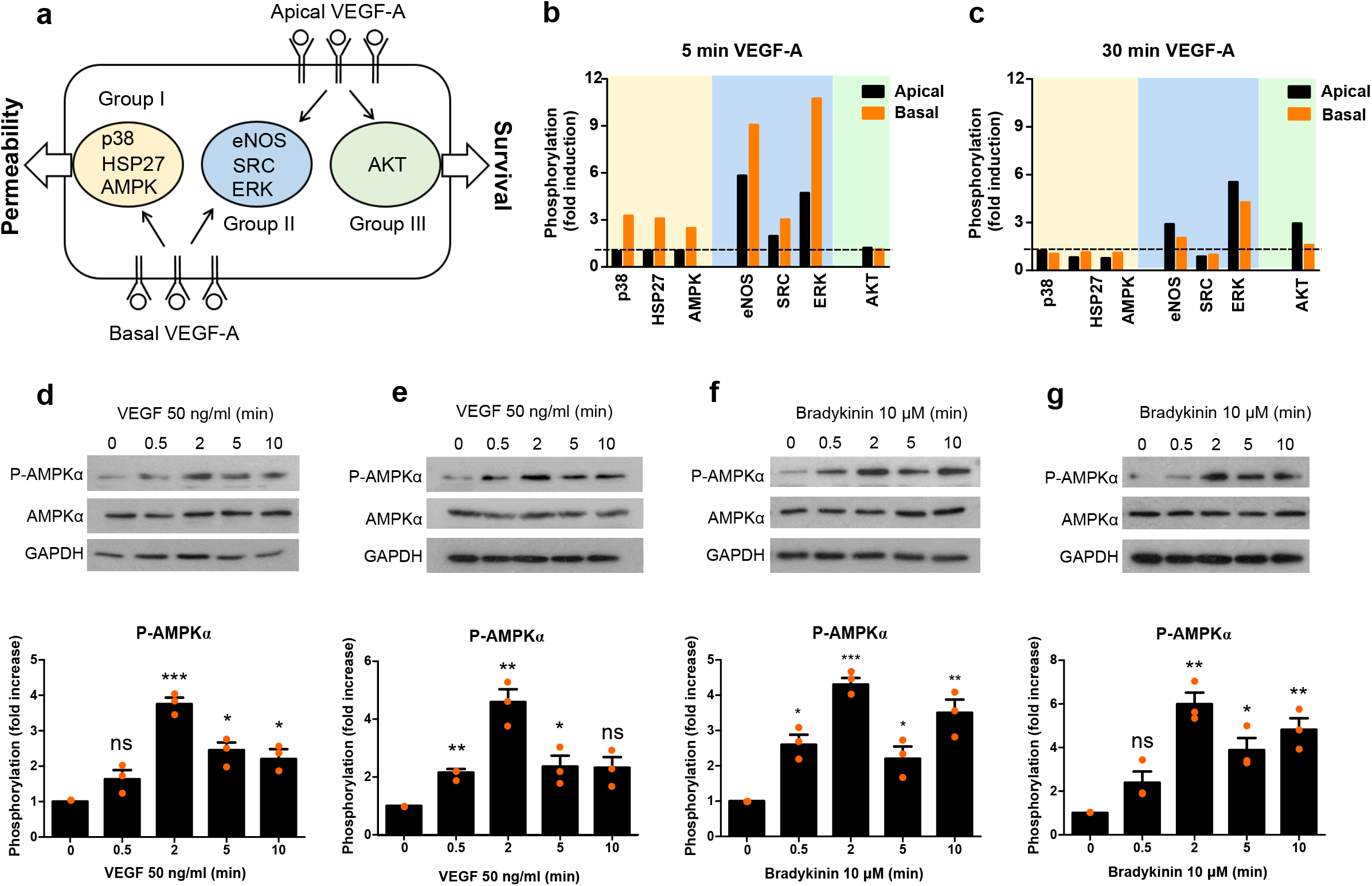
VEGF-A and Bradykinin induced AMPK phosphorylation. **(a-c)** Primary brain microvascular ECs were stimulated with VEGF-A from the apical or basal side for 5 min triggering apically (group I), basally specific (group III) or mixed (group II) responses (see text or ref 12 for more details) (a). Cells were lysed and phosphorylation of indicated molecules assessed by phosphoantibody array analysis (b-c). **(d-g)** brain microvascular ECs (d, f) or ex-vivo rat retinae (e, g) were stimulated with 50 ng/ml VEGF-A (d, e) or 10 μM BK (f, g) for the indicated length of time and AMPKα phosphorylation (pT172) analysed. Representative results and quantification of AMPKα activation from three independent experiments are shown as normalised means ± SD. *p < 0.05, **p < 0.01, ***p < 0.001.

Differentially phosphorylated proteins were categorised into three groups; group I phosphorylated only after basal stimulation with VEGF-A (such as p38, HSP27 and AMPK), group II phosphorylated regardless of the side of the stimulation (such as eNOS, SRC and ERK) and group III phosphorylated only when VEGF-A was applied apically (such as AKT). Phosphorylation of proteins exclusively in response to basally applied VEGF-A suggested they played a role in hyperpermeability (group I). Among these, AMPK stood out as its role in acute endothelial hyperpermeability has not yet been studied in detail. Additionally, AMPK was also among the 10 proteins, for which phosphorylation increased the most in response to basally applied VEGF-A, when analysed by phospho-peptide mass spectrometry (Dragoni and Turowski, unpublished results). A previous study also demonstrates that AMPK link Ca^2+^ transients to VE-cad phosphorylation in response to ICAM-1 engagement in brain microvascular ECs^26^.

AMPK phosphorylation in response to VEGF-A was confirmed by Western blot analysis. Stimulation of primary rat brain ECs from the basal side led to rapid, transient phosphorylation of AMPKα on T172, which peaked after ca. 2 min (Figure 3d). A very similar activation pattern was observed in the intact rat retina when VEGF-A was applied directly to the top of the isolated retina (corresponding to the basal side of the endothelium) (Figure 3e). BK induced similar phosphorylation of AMPKα T172, both in the cultured primary brain microvascular ECs and intact retina (Figure 3f, g). It was notable that AMPK phosphorylation was maximal after around 2 min in response to both VEGF-A and BK. However, BK clearly induced more sustained phosphorylation.

### AMPK mediates VEGF-A/Bradykinin-induced permeability

In order to specify the role of AMPK during VEGF-A- and BK-induced vascular permeability, AMPK activity was neutralised in the ex vivo retina. Preincubation of the ex-vivo retina with compound C, a widely used AMPK antagonist, significantly decreased VEGF-A- and BK-induced permeability by 80% and 93%, respectively (Figure 4a-d). Moreover, whole mount staining showed that pre-treatment with compound C prevented the VEGF-A- and BK-induced activation of p38, HSP27 and eNOS (Figure 4e, f).

**Figure 4.**
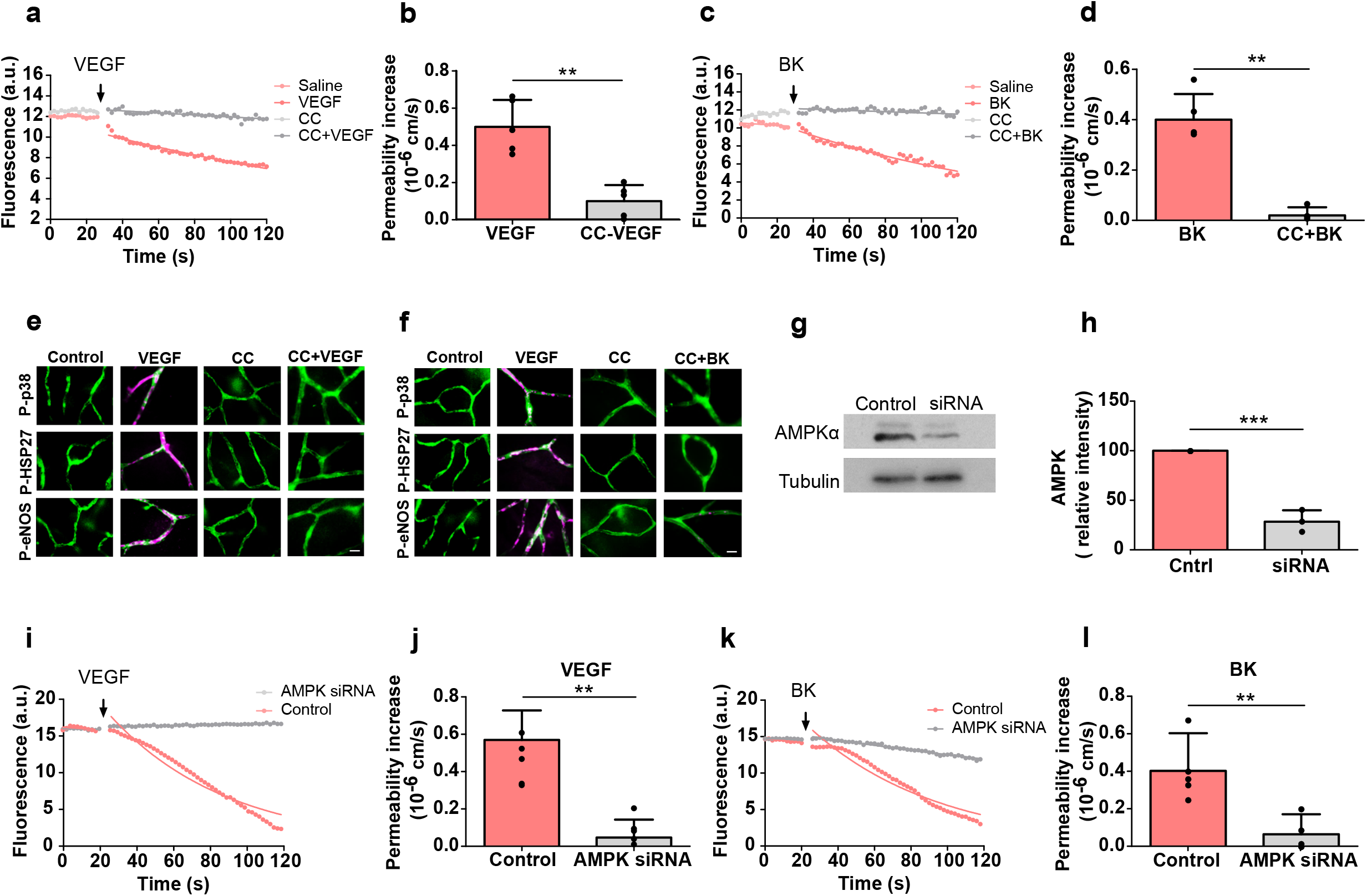
AMPK mediated VEGF-A/Bradykinin-induced permeability. **(a-f)** Rat ex-vivo retinae were preincubated with or without Compound-C (CC, 10 μM) for 15 minutes. 10 ng/ml VEGF-A (a, b) or 10 μM BK (c, d) were then applied to the top of the retina and changes in permeability were recorded. Alternatively (e, f), retinae were immunostained using IB4 (green) and anti-phospho-p38, -HSP27 or -eNOS (magenta) as detailed in Figure 2. **(g-l)** AMPKα specific siRNA or scrambled control was injected into the vitreous of mouse eyes. After 48 h retinae were isolated, lysed and subjected to immunoblotting as indicated (g). Shown in (h) is the densitometric quantification of 3 independent experiments as shown in (g). Alternatively, after 48 h retinae were prepared for ex vivo permeability measurements and stimulated using VEGF-A (50 ng/ml) (i, j) or BK (10 μM) (k, l). Note that neither PIF induced any permeability in the knocked down ex-vivo retina. Representative results and quantifications (normalized mean ± SD) from three independent experiments are shown. **p < 0.01, ***p < 0.001. Scale bars, 10 μm.

Next AMPKα was knocked down specifically by intravitreous injection of siRNA 72 h prior to preparing the retinae for ex vivo permeability measurements. Western blot results showed that AMPKα protein expression was significantly reduced by 70% (Figures 4g, h). Neither VEGF-A nor BK stimulation led to significant permeability in AMPK knocked-down retinae (Figures 4i-l).

Finally, when AMPK was directly activated by treating ex-vivo retinae with A769662 or AICAR, specific AMPK agonists that directly bind to and activate AMPK without any significant change in cellular ATP, ADP or AMP levels, baseline permeability was induced (Figure 5a-d) as well as the phosphorylation of AMPK, p38, HSP27 and eNOS (Figure 5e). Taken together these results confirmed the central role of AMPK in VEGF-A- and BK-induced acute retinal leakage and showed that AMPK acted upstream of p38 and eNOS.

**Figure 5.**
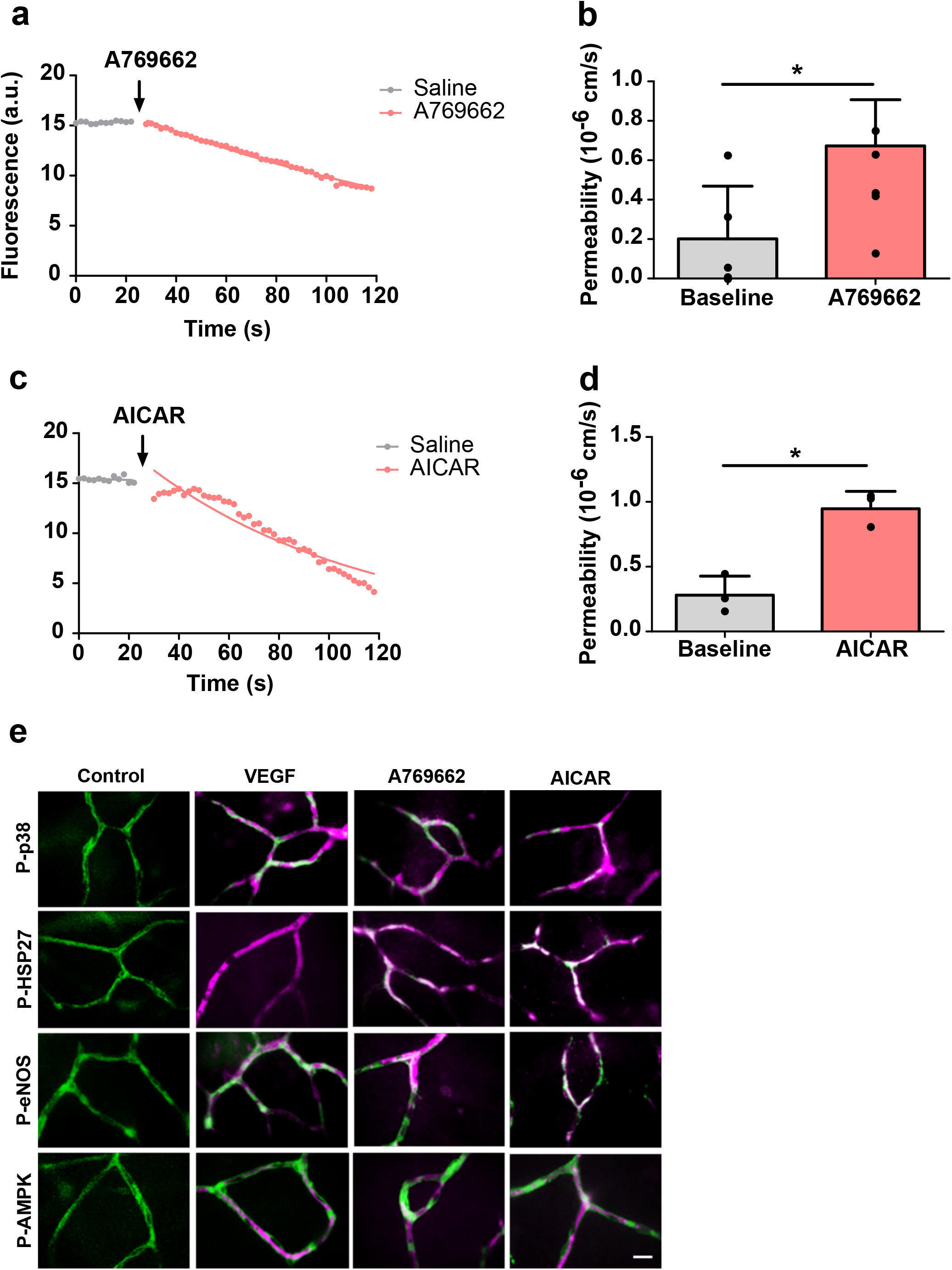
Stimulation of AMPK induced permeability in the ex vivo retina. **(a-d)** Ex vivo preparations were stimulated with the two different AMPK activators A769662 (10 μM) and AICAR (10 μM). Both agonists induced strong and immediate permeability in the ex vivo retinal microvessels. Mean (± SD) permeability changes recorded from at least four retinae are shown in (b, d). **(e)** Ex vivo retinae were stimulated as in (a-d) and after 2 min fixed using 4% PFA and then immunostained using IB4 (green) and or anti-phospho-p38, -HSP27 or -eNOS (magenta) as detailed in Figure 2 (magenta). *p < 0.05. Scale bars, 10 μm.

### VEGF-A- and BK-induced permeability requires Ca^2+^ and CAMKK, p38 and eNOS

Next, we aimed at placing AMPK within established PIF signalling cascades. Ca^2+^ is critical for the activation of both p38 and eNOS^26, 27, 28^. To test its role in VEGF-A-induced vascular leakage the ex-vivo retina was incubated with BAPTA, a Ca^2+^chelant, prior to VEGF-A administration. BAPTA treatment significantly reduced VEGF-A-induced permeability by 94%, and also prevented the phosphorylation of p38, Hsp27 and eNOS (Figures 6a, e), suggesting that Ca^2+^ acted upstream to these molecules. CAMKK is able to phosphorylate and activate AMPK in a Ca^2+^-dependent manner^26, 29^. Treatment of retinae with the CAMKK inhibitor STO-609 significantly reduced VEGF-A-induced permeability by 86% and also prevented the activation of p38, Hsp27 and eNOS (Figure 6b, e). The role of eNOS in permeability was re-confirmed by preincubation of the ex-vivo retinae with L-NAME, which reduced VEGF-A-induced permeability by 87% but it did not have any effect on p38 or HSP27 phosphorylation (Figure 6c, e), indicating that eNOS was not upstream of p38. Finally, the ex-vivo retina was preincubated with the p38 inhibitor SB203580, which significantly reduced the VEGF-A-induced permeability by 80% as well as activation of Hsp27 but not of eNOS (Figure 6d, e).

**Figure 6.**
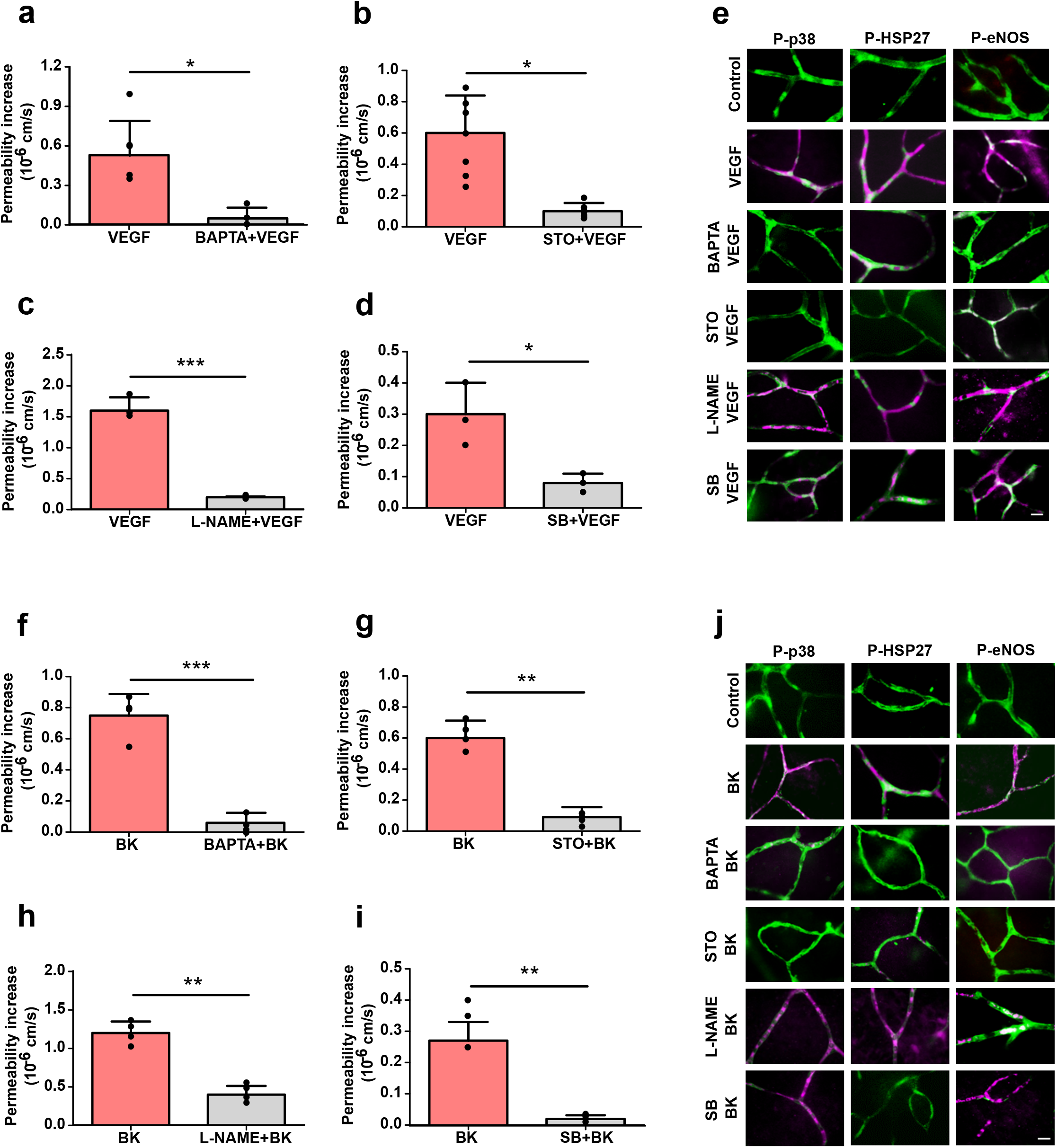
VEGF-A- and Bradykinin-induced permeability requires Ca2+, CaMKK, p38 and eNOS. **(a-e)** Ex vivo retinae were preincubated with 20 μM BAPTA (Ca^2+^ chelator) (a, e as indicated), 10 μM STO-609 (STO, CaMKK inhibitor) (b, e as indicated), 10 μM L-NAME (NOS inhibitor) (c, e as indicated), or 10 μM SB202190 (SB, p38 inhibitor) (d, e as indicated) for 15 minutes. Then VEGF-A (50 ng/ml) was applied to the top of the retina (abluminal side) and changes in microvessel permeability were recorded as described in Figure 2. Alternatively (e), retinae were fixed after 2 min using 4 % PFA and then immunostained with IB4 (green) and for phosphorylation of p38, HSP27 and eNOS (magenta). **(f-j)** As in panels (a-e), except that ex vivo retinae were stimulated with BK (10 μM). Representative results and quantifications (mean ± SD) from four independent experiments are shown. *p < 0.05, **p < 0.01, ***p < 0.001. Scale bars, 10 μm.

Similar results were obtained for BK-induced stimulation of retinae. BAPTA or STO nearly completely abolished BK-induced permeability, together with the activation of p38, Hsp27 and eNOS (Figure 6f, g, j). Pre-treatment of the ex-vivo retina with L-NAME reduced BK-induced permeability by 67% but did not affect p38 or HSP27 phosphorylation (Figure 6h, j). Finally, SB203580 prevented BK-induced permeability as well as phosphorylation of HSP27 but not eNOS (Figure 6i, j).

Importantly, VEGF-A- or BK-induced phosphorylation of AMPKα was completely abolished by BAPTA or STO. VEGF-A-but not BK-induced AMPKα phosphorylation was also abolished by SU1498 (Figure 7a). These results indicated that in the retina, VEGF-A and BK induced Ca^2+^ transients and consequent activation of CAMKK and AMPK. At this point signalling diverged into either activation of p38 or eNOS, which both contributed to permeability (Figure 7c).

**Figure 7.**
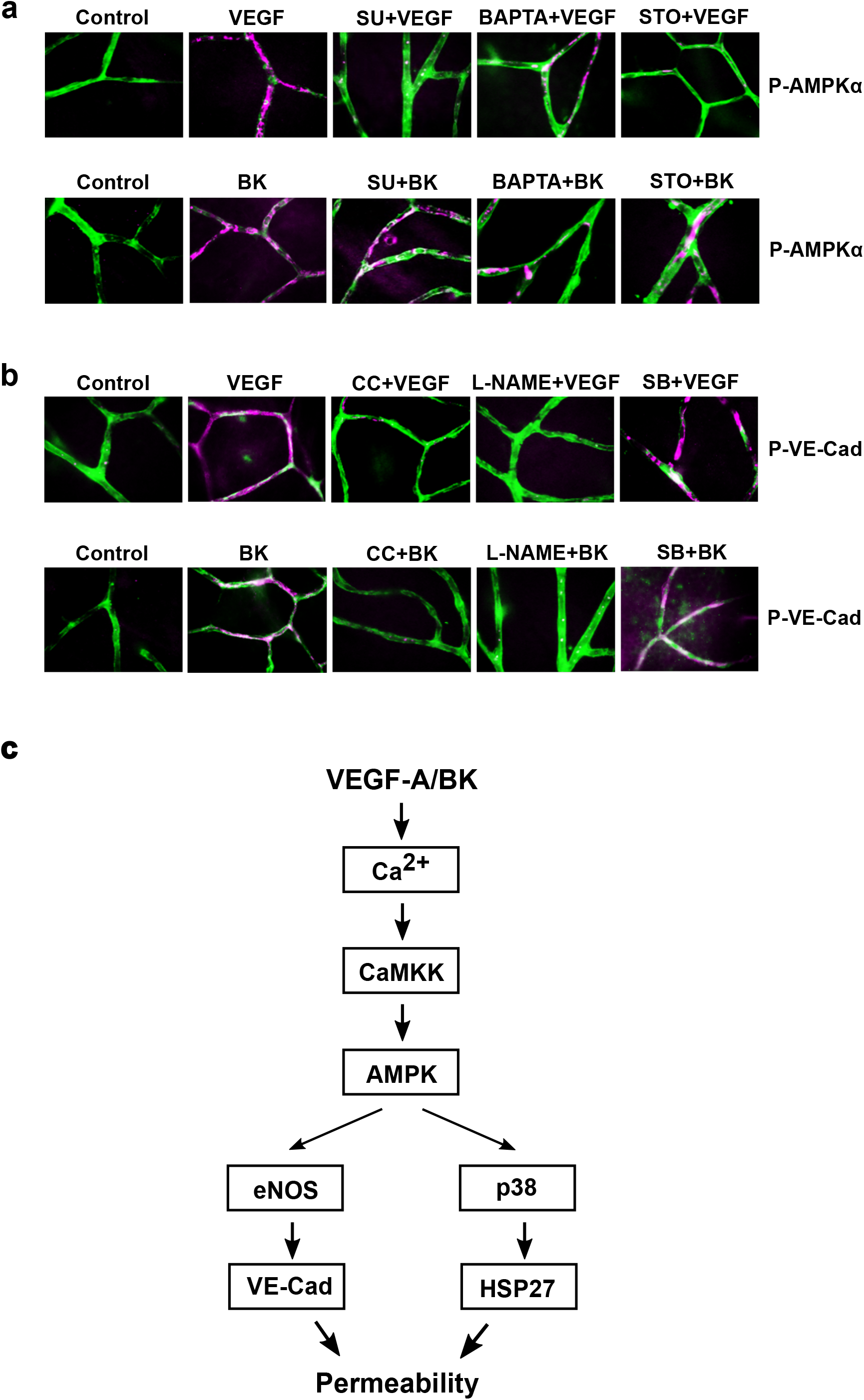
VEGF-A and Bradykinin induced VE-Cadherin phosphorylation. **(a)** Ex vivo retinae were preincubated with SU1498 (10 μM), or BAPTA (20 μM) or STO (10 μM) and treated with VEGF-A and Bradykinin for 2 min. Retinae were then fixed with 4% PFA and immunostained for phospho-AMPKα (magenta) and with IB4 (green). **(b)** Ex vivo retinae were preincubated with compound C (10 μM), or L-NAME (10 μM) or SB202190 (10 μM), treated with VEGF-A and Bradykinin for 2 min and immunostained for phosphorylated VE-cad (using anti-pY685-VEC) (magenta) and with IB4 (green). **(c)** Proposed signalling networks in the ex-vivo retinal microvasculature downstream of VEGF-A and Bradykinin. Scale bars, 10 μm.

### VEGF-A and BK stimulate VE-Cadherin phosphorylation

Lastly, we wanted to analyse VE-cad internalisation in the VEGF-A and BK stimulated ex vivo retina. However, systematic correlation of cryo-immuno EM to leaky microvessels proved impractical. We therefore used phosphorylation of VE-cad on Y685 as a surrogate marker for retinal microvessel permeability^16^, in particular since such VE-cad phosphorylation is also a prerequisite for internalisation after BK treatment^15^. VEGF-A or BK stimulation of ex vivo retinae induced tyrosine phosphorylation of VE-cad on Y685 in intact microvessels (Figure 7b). This phosphorylation was completely abolished following pre-incubation with compound C, L-NAME but not SB203580, indicating that VE-cad was downstream of AMPK/eNOS but not AMPK/p38 (Figure 7c).

## DISCUSSION

Measuring retinal microvascular permeability has been mainly restricted to Miles-type assays using either fluorescent tracers or Evans Blue/albumin^8, 30^. Chronic leakage can also be visualised in real time by fluorescein angiography^31^. However, collectively these methods, which measure the amount of extravasated tracer, do not only reflect the degree of leakage but are also strongly influenced by dye concentration in the vasculature and dye clearance from the tissue^1, 2^. Further disadvantages of these methods are that compound concentrations and timing cannot be controlled accurately. Thus, they are inadequate to measure acute permeability and associated signalling accurately and in a controlled manner.

The ex-vivo retinal platform described herein addressed most of these issues. It constituted a significant advance to EC cultures, since it used a complete and intact neurovascular unit. The functionality of the retinal vasculature was preserved, with both VE-cad and claudin-5 distribution indistinguishable from that in vivo. Similar preparations of the brain and the retina also display full cellular functionality of e.g. pericytes^32^. Importantly, permeability to sulforhodamine-B was very low and within the range of that of other small non-ionic molecules at the intact BBB in vivo^33^ and notably ca. 10 x lower than in preparations of pial microvessels^34^, indicating that this model was highly suitable for permeability measurement at an intact neurovascular unit. Vascular barrier properties in these retinal explants was, as expected, dependent on tight junction integrity. The reported speed, at which compounds such as VEGF-A induce permeability^10^, was fully recapitulated. Leakage measurements were then combined with whole tissue staining and analyses of the phosphorylation status of key mediators of permeability using phospho-specific antibodies to gain mechanistic insight. Furthermore, the preparations could be interrogated using small molecule antagonists and agonists at defined concentrations and times, and allowed for the identification of key downstream regulators, common to both VEGF-A and BK stimulation. Even more specific neutralisation of key proteins was achieved through prior intraocular injection of siRNA. Conceivably this model is compatible for use with genetically modified mice and disease models, further broadening its applicability. The ex vivo platform could also be used to measure Ca^2+^ transients or localised production of reactive oxygen or nitrogen species in response to vasoactive compounds such as VEGF-A or BK.

Measurements in retinae were done ex vivo, in the absence of blood flow. Blood flow and associated shear stress may influence EC biology, such as cell-cell adhesion and inflammatory dysfunction^15, 35, 36^ and their absence in our preparation must be taken into account when evaluating results. However, it should also be noted that permeability regulation by shear stress appears to be remembered in ECs in vitro for at least 24 h^37^.

For the purpose of developing and validating the ex vivo retinal platform we have focused on PIFs, which induce permeability when added to the abluminal (tissue) side of the endothelium, such as VEGF-A and BK^12^, as these are readily applied on top of the retinal explants. Other PIFs that act only from the luminal side, such as lysophosphatidic acid^12^ or lysophosphatidylcholine^8^, could also potentially be investigated in this system. However, this would require the use of a manifold injection system^38^, which allows switching between injection of sulforhodamine-B with or without permeability factor into radial vessels.

VEGF-A and BK induced acute leakage in retinal microvessels, which was associated with and dependent on Ca^2+^, the MAPK p38 and eNOS, in agreement with published data^11, 12, 14^. We also identified AMPK as a novel key mediator of both VEGF-A- and BK-induced permeability, indicating it is a core regulator of acute vascular permeability. AMPK is primarily known to regulate energy requirements of the cell, but has also been implicated in other seemingly unrelated cellular processes such as migration, cell growth and apoptosis^39^. This protein kinase has been studied before in relation to its protective role of the BBB^19, 20^, whereas in the retinal pigment epithelium it has been shown to be responsible for the permeability induced by IL-1β^21^. However, all these studies address chronic changes and do not focus on the role of AMPK for acute permeability. Whilst its canonical activation is dependent on cellular AMP: ATP concentrations and phosphorylation on T172 by LKB1, we found that, in the regulation of endothelial permeability, AMPK was activated downstream of Ca^2+^ and CAMKK. This activation pathway has previously been described as non-canonical^40^ and is also operational when CNS ECs facilitate the transmigration of lymphocytes^26^. Notably, VEGF-A has also been reported before to induce NO production via a pathway requiring Ca^2+^ and AMPK^41^. Indeed, eNOS phosphorylation on S1177 can be mediated by AKT^42^ or AMPK^26, 43^. However, the PI3K/Akt pathway is not relevant to VEGF-A-induced permeability induction in neurovascular ECs^12^. We confirmed the phosphorylation of eNOS on S1177 downstream of AMPK both during VEGF-A and BK permeability induction. Interestingly, the Ca^2+^/AMPK/eNOS pathway resulted in the phosphorylation of VE-cad on Y685, identified previously as key for vascular permeability in the periphery^15^ and the retina^16^, thus providing a direct link between AMPK and paracellular junction regulation. For retinal permeability, occludin phosphorylation and internalisation also plays an important role. However, judging from the published time courses, it is likely to be a later event, not captured by our experiments, and thus either secondary to VE-cad phosphorylation or, with its dependency on PKCß activation, outside of the signalling network we have investigated here^17^.

In response to VEGF-A and BK, AMPK also regulated the phosphorylation of the MAPK p38 and its substrate HSP27, both previously implicated in actin rearrangement during endothelial barrier disruption^44, 45^. P38 is a bona fide regulator of VEGF-A responses^46^ and its activation downstream of cdc42 and PAK and subsequent modulation of the actin cytoskeleton occurs during VEGF-A-induced endothelial migration downstream of VEGFR2 phosphorylation on Y1214^47^. Whilst it is possible that additional, parallel AMPK regulation of this cascade was operational during VEGF-A or BK-induced permeability, we favour an alternative model of direct activation of p38 by AMPK via TAB1, a pathway described in apoptotic lymphocytes and the ischemic heart^48, 49^. By switching between two different p38 activation modes (Ca^2+^/AMPK/TAB1 versus cdc42/PAK) ECs could adapt cytoskeletal regulation to the specific requirement of EC migration or permeability.

The ex-vivo retina proved to be a reliable model and demonstrated its usefulness in identifying key regulators of acute permeability. Whilst AMPK clearly emerged as such a key regulator, it is unlikely to be exploitable as a target for anti-leakage treatments: its central role in regulating cellular energy demands throughout the body hints at many potential side effects. Activation of AMPK is currently investigated as a therapeutic option to treat cancer, metabolic syndrome and diabetes^50, 51^. However, in light of the strong induction of permeability we observed in response to at least two AMPK agonists, we propose that these avenues should be explored cautiously, since at least acute microvascular leakage may accompany such treatment modalities. Nevertheless, our data collectively indicated that the ex vivo retina platform can play an important part in elucidating mechanisms and signalling of neurovascular leakage.

## ACKNOWLEDGEMENTS

We thank Dr Paul Fraser (King’s College London) for introducing us to his early ex-vivo retinal preparations and Prof. John Greenwood (UCL) for critical comments on the manuscript. siRNAs targeting CLDN5 were kindly provided by Dr Matthew Campbell (Trinity College Dublin). Phospho-VE-Cad antibodies were kindly provided by Dr Fabrizio Orsenigo and Prof. Elisabetta Dejana (FIRC Institute of Molecular Oncology, Milan and Uppsala University).

This work was supported by a Moorfields Eye Charity grant to S.D. and a Diabetes UK grant to P.T.

The data that support the findings of this study are available from the corresponding author upon reasonable request.

Supplemental material for this paper can be found at the journal website: https://eur01.safelinks.protection.outlook.com/?url=http%3A%2F%2Fjournals.sagepub.com%2Fhome%2Fjcb&data=02%7C01%7C%7Cc27df537f41f419d715808d795e9c351%7C1faf88fea9984c5b93c9210a11d9a5c2%7C0%7C0%7C637142703553285613&sdata=J8jD0hoMtCdGE1jbrGR5QDN1QGWyibw0hnEBOenTqj0%3D&reserved=0

## AUTHOR CONTRIBUTION STATEMENT

S.D.: designed experiments; performed experiments; analysed the data; wrote the manuscript

B.C.; E.K.; T.B; M.H.S.: performed experiments

P.T.: designed experiments; performed experiments; analysed the data; wrote the manuscript

---

**Supplemental Figure 1.**
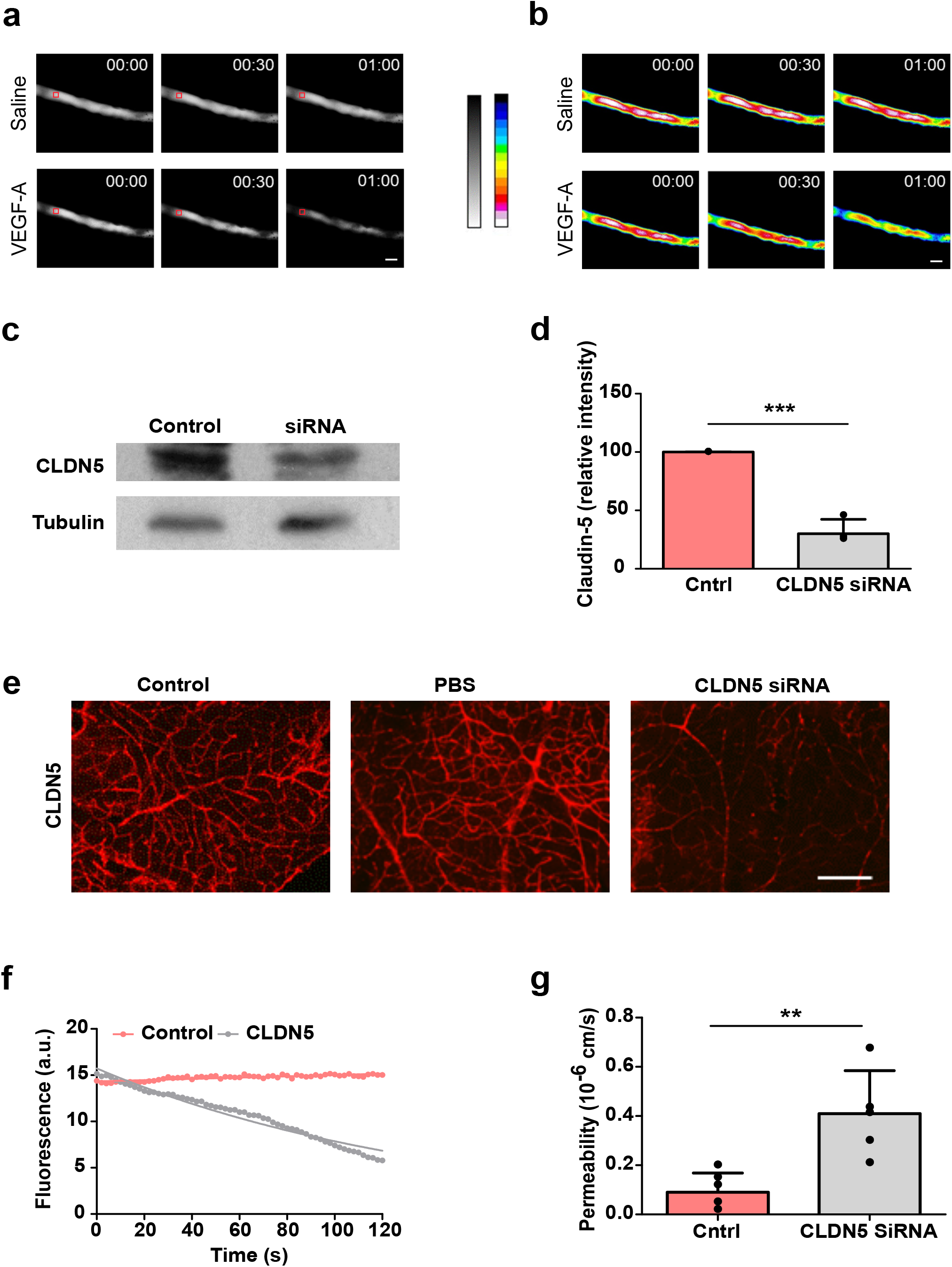
Properties of the ex vivo retina model. **(a-b)** Selected frames, pseudocoloured in (b), of time-resolved recordings of a Sulforhodamine-B-filled rat capillary before and after the addition of 50 ng/ml VEGF-A, illustrating the rapid loss of fluorophore from the lumen of the vessel. The red box exemplifies a typical r.o.i used for intensity measurement. Scale bars, 10 μm. **(c-g)** CLDN5 siRNA was injected into mouse eyes. 48 h later CLDN5 levels were analysed by immunoblots of retinal lysates (c-d) or by wholemount immunochemical staining (e). Scale bar, 100 μm. Alternatively, permeability of 4 KDa Rhodamine was measured in ex vivo retinae from CLDN5 siRNA or control injected eyes (f-g). **p < 0.01, ***p < 0.001.

